# Quantifying Leaf Symptoms of Sorghum Charcoal Rot in Images of Field-Grown Plants Using Deep Neural Networks

**DOI:** 10.1101/2024.04.17.589978

**Authors:** Emmanuel Gonzalez, Ariyan Zarei, Sebastian Calleja, Clay Christenson, Bruno Rozzi, Jeffrey Demieville, Jiahuai Hu, Andrea L. Eveland, Brian Dilkes, Kobus Barnard, Eric Lyons, Duke Pauli

**Affiliations:** School of Plant Sciences, University of Arizona, Tucson, AZ; Department of Computer Science, University of Arizona, Tucson, AZ; Donald Danforth Plant Science Center, St. Louis, MO; Department of Biochemistry, Purdue University, West Lafayette, IN; Center for Agroecosystem Research in the Desert (ARID), Tucson, AZ

**Keywords:** Sorghum, Charcoal Rot of Sorghum, Plant Disease, Machine Learning, Image Analysis

## Abstract

Charcoal rot of sorghum (CRS) is a significant disease affecting sorghum crops, with limited genetic resistance available. The causative agent, *Macrophomina phaseolina* (Tassi) Goid, is a highly destructive fungal pathogen that targets over 500 plant species globally, including essential staple crops. Utilizing field image data for precise detection and quantification of CRS could greatly assist in the prompt identification and management of affected fields and thereby reduce yield losses. The objective of this work was to implement various machine learning algorithms to evaluate their ability to accurately detect and quantify CRS in red-green-blue (RGB) images of sorghum plants exhibiting symptoms of infection. EfficientNet-B3 and a fully convolutional network (FCN) emerged as the top-performing models for image classification and segmentation tasks, respectively. Among the classification models evaluated, EfficientNet-B3 demonstrated superior performance, achieving an accuracy of 86.97%, a recall rate of 0.71, and an F1 score of 0.73. Of the segmentation models tested, FCN proved to be the most effective, exhibiting a validation accuracy of 97.76%, a recall rate of 0.68, and an F1 score of 0.66. As the size of the image patches increased, both models’ validation scores increased linearly, and their processing time decreased exponentially. The models, in addition to being immediately useful for breeders and growers of sorghum, advance the domain of automated plant phenotyping and may serve as a base for drone-based or other automated field phenotyping efforts. Additionally, the models presented herein can be accessed through a web-based application where users can easily analyze their own images.

**Core ideas:**

1. Automated phenotyping tools are required for the efficient detection and quantification of charcoal rot of sorghum.
2. Classification and segmentation models can distinguish between concurrent plant stresses with similar symptoms.
3. Larger image patch sizes generally improve model performance and reduce processing time.

## 1 INTRODUCTION

The impacts of plant biotic stresses pose a significant risk to sustainable agricultural production and threaten the availability of nutritious calories to a growing world population. Globally, plant diseases are directly responsible for yield losses ranging from 10 to 40% across major staple crops that provide approximately 50% of the calorie intake among humans (Savary et al. 2019). Further compounding cropping system challenges created by biotic stressors is the presence and interaction with abiotic stresses such as heat and drought (Desaint et al. 2021; Pandey, Ramegowda, and Senthil-Kumar 2015; H. Zhang and Sonnewald 2017; Ramegowda and Senthil-Kumar 2015). Collectively, these factors further exacerbate the challenges facing crop production and highlight the need for resistant crop cultivars capable of mitigating these factors (Kissoudis et al. 2014). This urgent need for improved cultivars is further highlighted by the aridification of agricultural lands which will increase and intensify the effects of abiotic and biotic stresses (Overpeck and Udall 2020); these environmental changes will likely disrupt and alter the geographic distribution and abundance of crop pathogens (Newbery, Qi, and Fitt 2016; Chakraborty, Tiedemann, and Teng 2000; Delgado-Baquerizo et al. 2020).

Traditional plant breeding methods have relied on the visual assessment and scoring of germplasm subjected to infection, either naturally or artificially inoculated, to identify genotypes with varying levels of disease resistance (St Clair 2010; Bernardo 2014). However, visual assessment by trained experts is subject to human biases and errors that reduce the precision, accuracy, and repeatability of disease rating resulting in decreased selection accuracy, heritability, and genetic gain (Poland and Nelson 2011; Bock et al. 2009), which, in turn, lengthens the development of cultivars that can cope with biotic and abiotic stresses. To overcome these limitations, machine learning (ML) and artificial intelligence (AI) algorithms, in conjunction with high-throughput phenotyping, can be leveraged to conduct automated and rapid assessment of images of diseased plants/tissue to provide more accurate and reliable scoring of relevant plant germplasm (DeChant et al. 2017; Pauli et al. 2016; Singh et al. 2016; Lu, Tan, and Jiang 2021). In turn, these data can be used for applied breeding either through direct selection methods or by identifying and characterizing genes or alleles that confer resistance to a pathogen. These methods can also be used by growers for the early and rapid detection of disease and plant stress, allowing for more timely mitigation strategies.

*Macrophomina phaseolina* (Tassi) Goid is a pluriparous plant pathogen impacting over 500 plant species in more than 100 plant families, including cereals, legumes, vegetables, and fruits throughout the world (Kunwar et al., n.d.; Marquez et al. 2021). *M. phaseolina* is a necrotrophic soilborne fungus native to the Sonoran Desert soil (Mihail et al. 1989; Mihail, Ramussen, and Turner, n.d.) but is also widely distributed in the United States. It survives and spreads primarily as black microsclerotia in diseased root and stem debris as well as in soil after decay of infected plant material (Bhattacharya and Samaddar 1976). Microsclerotia infection of root tissue occurs at temperatures ranging from 20°C to 40°C, affecting plants at different developmental stages, including seedling, young, and mature phases (Hsi, n.d.; Collins, Wyllie, and Anderson 1991). Disease development is influenced largely by drought stress and low soil moisture (Odvody and Dunkle, n.d.). Under favorable environmental conditions, *M. phaseolina* invades the vascular system, disrupting the normal function of water and nutrient transport to the leaves, causing visible symptoms such as wilting and premature leaf death. In general, the onset of symptoms varies with host species and cultivars, plant growth stage at the infection, and environmental conditions.

Seedlings can be infected at emergence through the early vegetative stages, but symptoms are not typically observed until plants undergo drought stress or physiological stress at later growth stages (pollination or grain filling). Early foliar symptoms include wilting leaves with grayish white appearance (e.g., frost-damaged), reduced vigor, and premature plant death scattered individually or in small patches throughout a field. Unfortunately, these foliar symptoms are not unique to *M. phaseolina* and often are confused with those of other disorders and diseases such as drought stress, frost-damage, early senescence, root rot, and Fusarium rot.

Sorghum [*Sorghum bicolor (L.)* Moench] is a key cereal crop that also serves as a host species for *M. phaseolina*. Globally, sorghum is the fifth most widely grown cereal crop, trailing maize, rice, wheat, and barley, and is a staple food crop for millions living in semi-arid regions (Hossain et al. 2022). In the US, sorghum is gaining popularity given its innate ability to produce grain on marginal land and areas that lack sufficient irrigation water needed for row crops such as maize, and because of its diverse utility as a food, feed, and biofuel crop (Yang et al. 2022; Ndlovu, van Staden, and Maphosa 2021; Tang et al. 2018; Rai et al. 2016; Rooney et al. 2007). The range of sorghum’s end uses combined with its innate genotypic and phenotypic diversity are helping to position this crop as a sustainable solution to agricultural production in the face of climate change (Boatwright et al. 2022; Chadalavada, Kumari, and Kumar 2021). However, for sorghum production acreage to increase, resistance to charcoal rot of sorghum (CRS, *M. phaseolina* infection in sorghum) is needed as CRS causes a variety of symptoms including root rot, soft stalk, early lodging of plants, premature drying of stalk, reduced head size, and poor filling of grain (Hsi, n.d.). These symptoms and agronomic impacts greatly reduce the performance and hence profitability of the crop. Further compounding the problems associated with CRS, is that typical sorghum production occurs in arid and semi-arid environments, regions prone to water deficit stress, increasing the co-occurrence of both biotic and abiotic stress that negatively impact yield. The manifestation of both types of stress in the growing environment results in similar visible phenotypes, making discriminating between the two quite challenging.

Given the visual similarity between sorghum leaves impacted by CRS and drought stress (ie. water deficit), an automated phenotyping tool that accurately identifies and quantifies the foliar symptoms of CRS versus drought stress would be valuable for researchers, breeders, and crop managers in arid production regions. Furthermore, an image-based phenotyping solution that can be deployed on small, unoccupied aerial vehicles (UAVs) or handheld devices such as smartphones would further increase the utility of such a tool. To accomplish this, it will be crucial to utilize advancements in both hardware and software to create “lightweight” algorithms. These can be implemented on edge devices, allowing computation and processing to occur closer to the data source, right in the field. With the advent of powerful computation devices, specifically graphical processing units (GPU), and recent advancements in ML and AI algorithms, many domain scientists have started to use neural networks (NNs) to detect, locate and quantify many features, including disease, in different modality of images (Amsaveni and Albert Singh 2013; Mechria, Gouider, and Hassine 2019; Siar and Teshnehlab 2019). The class of NNs that are widely applied for this application are known as convolutional neural networks (CNNs) (LeCun 2015). By learning useful features from image data automatically, these CNNs can perform classification or detection tasks more effectively than the older approach of manual feature extraction (Rybski et al. 2010). The CNNs have also been used for detection, classification, and segmentation of similar foliar diseases in other crop species. In DeChant et al. (2017), the authors proposed a computational pipeline of CNNs for classifying images of field-grown maize images to determine presence or absence of Maize Northern Leaf Blight (NLB). Expanding on this work, Stewart et al. (2019) trained a mask region-based convolutional neural network (mask R-CNN) for segmenting maize images acquired by UAVs into healthy and NLB-affected tissues. In Wu et al. (2019), the authors proposed a sliding window approach for generating heat maps highlighting regions in the aerial images of maize fields affected by the NLB. Wu et al. (2019) and Weisner-Hanks et al. (2019) proposed a combination of CNNs and conditional random fields (CRF) to segment maize images into normal and NLB-affected regions. For early plant disease detection, hyperspectral imaging is being increasingly utilized as an alternative to RGB images, given its ability to offer unique spectral signatures and valuable insights into plant health (Yu et al. 2018; Mertens et al. 2021; G. Zhang et al. 2022). However, its complexity and high computational demands present challenges, hindering its full implementation in agriculture (Cheshkova 2022; Okyere et al. 2023). Meanwhile, standard RGB cameras simplify the process and broaden the application of disease detection methods from research to commercial agriculture.

In the present study, we propose two CNN-based approaches for quantifying CRS in RGB images captured of field-grown sorghum under drought stress conditions and compare the performance of the two methods. The first approach, which involves a set of classification models, determines the presence of CRS in images by classifying small-sized patches. The second approach carries out pixel-wise classification or semantic segmentation on the images. We evaluate the performance of these two approaches to determine which is more capable of detecting and quantifying CRS as well as provide computational benchmarks for deployment by end-users. We also provide a high-quality labeled dataset for classification and segmentation tasks as well as our code for the benefit of other researchers to improve upon our work. To reduce the barrier to entry, a web application is provided which allows end-users to deploy all models in the present study.

## 2 MATERIALS AND METHODS

### 2.1 Data and Field Experiments

A population of ethyl methanesulfonate (EMS)-mutagenized BTx623 sorghum, the genotype used for the generation of the sorghum reference genome, consisting of 430 individuals (Addo-Quaye et al. 2018; Paterson et al. 2009) was evaluated at the Maricopa Agricultural Center (MAC) of the University of Arizona located in Maricopa, AZ (33°04’37” N, 111°58’26” W, elevation 358 m) in 2020. The population was grown under contrasting irrigation conditions representing well-watered (WW) and water-limited (WL) conditions. The trial was planted on June 17 (day 169, Julian calendar) in a partially replicated, incomplete block design with 240 of the lines replicated within each of the two irrigation treatments while 190 lines were only observed once per irrigation treatment. The order of entries within each irrigation treatment was randomized. To reduce edge effects, the wildtype BTx623 sorghum cultivar was planted at the perimeter of each irrigation treatment. Experimental units were one-row plots, 3.5 m in length with a 0.5 m alley at the end of each plot and inter-row spacing of 0.76 m; plots were thinned to a density of five plants per plot (1 plant per 0.7 linear m) after crop establishment, at approximately V5 growth stage. The soil type is a Casa Grande sandy loam (fine-loamy, mixed, superactive, hyperthermic Typic Natrargids). Conventional sorghum cultivation practices (fertilizer application rate/amount, weed/insect control, etc.) for the desert Southwest were employed (Ottman 2016). Meteorological data were obtained from an automated Arizona Meteorological Network (AZMET) weather station (cals.arizona.edu/AZMET/06.htm) located on the premise of MAC and 738 m from the field (Brown 1989). Crop irrigation was performed using subsurface drip irrigation with pressure compensated drip tape (DripNet PC, Netafim, Tel Aviv, Israel) buried at a depth of approximately 0.15 m beneath the soil surface, directly underneath the plants. Soil volumetric water content (SVWC) was monitored on a biweekly basis using a field-calibrated neutron moisture probe (Model 503, Campbell Pacific Nuclear, CPN, Martinez, CA, USA) with measurements taken in 0.2 m increments from a depth of 0.1 m to 1.9 m so that irrigation amounts could be adjusted to achieve approximately 25% and 14% SVWC in the WW and WL treatments, respectively.

### 2.2 Disease Presence and Description

Symptomatic disease tissue was observed in the field trial approximately five weeks after planting with plants exhibiting typical signs of wilting and root rot (Figure 1). Initially, about 15% of research plots showed severe wilting symptoms; however, this slowly increased as the season progressed, and ultimately approximately 70% of the research plots exhibited some level of infection. The presence of similar symptoms was also confirmed previously in 2019 and 2017 within the same field as used for the present work. To confirm pathogen presence and identity, symptomatic plants were dug up from the field plots ensuring that roots remained attached to the plant while avoiding damage to the roots. All sampled plants were placed in plastic bags and kept cool until they could be transported to the University of Arizona’s Extension Plant Pathology lab in Tucson, AZ, USA.

**Figure 1.**
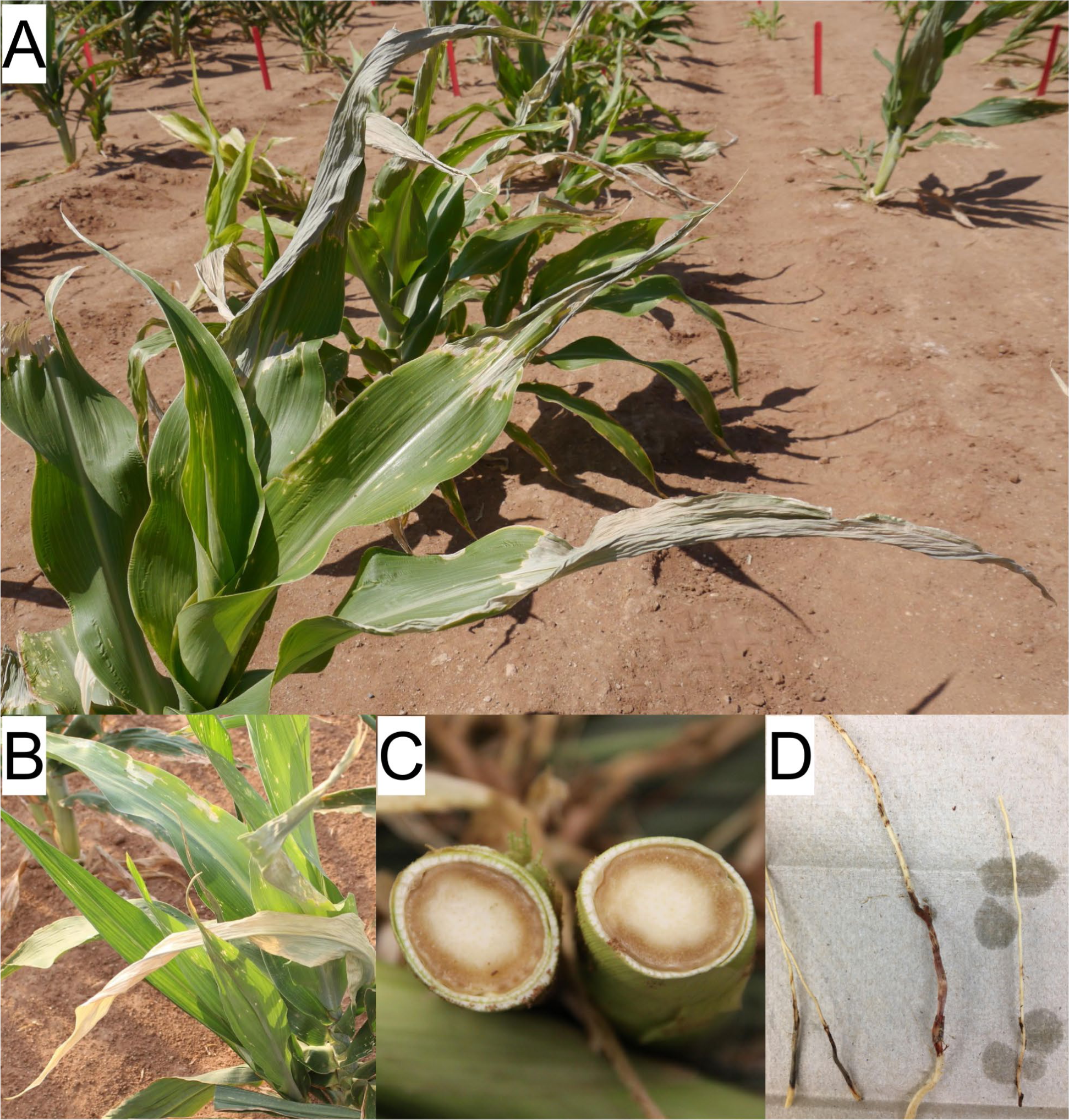
Symptoms of young sorghum plants infected with charcoal rot of sorghum (CRS), a fungal disease caused by *M. phaseolina*. A) Grayish white appearance of young plants in a field in Maricopa, Arizona. B) Close up of a young sorghum plant displaying symptoms of CRS, including leaf curling and hooking, chlorosis, and necrosis. C) Discoloration of stem vascular tissue. D) rotting of roots.

Since microsclerotia are not visible in the stem, isolation was made using putatively infected root and stem tissues (Figure 1C and D). Tissue samples, measuring 7 x 7 x 3 mm, were cut from the margin between the diseased and seemingly healthy tissues. These tissue samples were surface sterilized by soaking in 75% ethanol for five seconds, 1% sodium hypochlorite for one minute, rinsed well with sterile distilled water, and dried on sterile filter paper in a laminar hood. Each sterile tissue sample was plated onto potato dextrose agar (PDA) plates and water agar plates. Plates were incubated at 25°C in the dark until fungal colonies were observed. Colonies were subcultured onto PDA plates and incubated at 25°C for four days. Hyphal tip subcultures were obtained for each isolate from the colony margin and subcultured onto fresh PDA. Morphological characteristics were observed on 3 isolates on two-week-old PDA cultures. Based on the culture morphology on PDA (Figure 2), all three isolates were tentatively identified as *M. phaseolina*. To confirm its identity, genomic DNA was extracted from mycelial mats of three isolates using DNeasy Plant Pro Kit (Qiagen Inc., Valencia, CA) according to the manufacturer’s instructions. The internal transcribed spacer (ITS) region of the rRNA gene was amplified with primers ITS1/ITS4 and three nucleotide sequences were obtained. A BLASTN search revealed that sample sequences shared a 100% match with sequences of *M. phaseolina* in the NCBI GenBank Database (Sayers et al. 2022).

**Figure 2.**
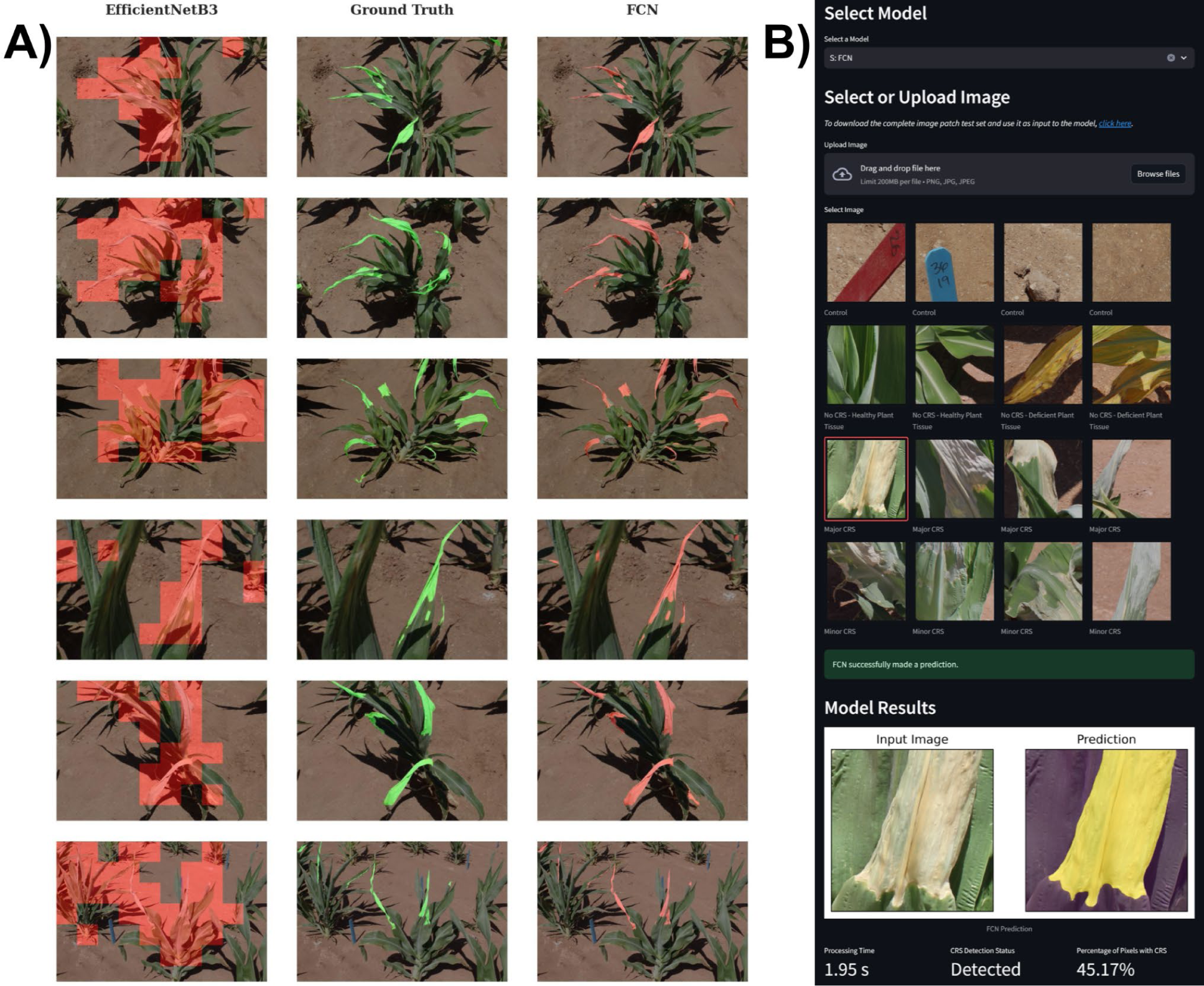
Comparison and deployment of classification and segmentation models. A) Comparison of the identification of Charcoal Rot of Sorghum (CRS) by the classification model EfficientNet-B3 and the segmentation model Fully Convolution Network (FCN). Ground truth, hand-labeled images are in the center column while the EfficientNet-B3 and FCN model outputs are on the left and right hand sides, respectively. For EfficientNet-B3, image patches of size 512 x 512 are highlighted since the model only classifies whether or not the image contains CRS-affected pixels. The FCN model, which performs segmentation, highlights the specific pixels that are affected with CRS closely resembling the results of the hand-labeled, ground truth data. Both models were trained on image patch sizes of 512 x 512 pixels. B) A Streamlit application for easy deployment of all classification and segmentation models presented here. The application allows users to (*i*) select a model, (*ii*) upload an image or select one from the image gallery of test set images, and (*iii*) obtain model results including processing time, detection status, and percentage of pixels with CRS. The application can be accessed at: https://charcoal-dryrot-quantification.streamlit.app/.

### 2.3 Plant Imaging

#### 2.3.1 Image Capture

Images of both infected and noninfected sorghum plants were taken 51 days after planting, equating to GS-1 vegetative (germination to panicle initiation) growth stage (Roozeboom and Prasad 2019). Images were collected from both the WW and WL irrigation treatments. A total of 1,400 high-resolution, JPEG-formatted images 5184 x 3456 and 2336 x 1752 pixels were taken of sorghum plants within the field using a Canon Rebel T6 camera (Canon, Tokyo, Japan) and a Sony Alpha a6000 (Sony, Tokyo, Japan), respectively. The camera operators walked through the field and images were taken at random to include a variety of angles, lighting conditions, background features, and zoom settings. The image capture settings of both cameras were set to auto adjust (aperture, shutter speed, and ISO speed) to ensure adequate and unbiased image capture given the number of images that had to be taken. All images were taken on the same day between 10:00 to 14:00. Collected images were uploaded to the CyVerse Data Store for further processing (Devisetty et al. 2016).

The 1,400 CRS-impacted sorghum images were imported into the image labeling platform Labelbox (https://labelbox.com/, verified on 2/7/2024) and annotated by researchers trained in CRS identification. Researchers were instructed to label the area exhibiting symptoms of CRS by drawing high-fidelity polygons around CRS-affected sorghum plant tissue. After annotation, images were reviewed and validated by a second set of experts and split into three different sets: training (60%, 840 images); validation (20%, 280 images); and test (20%, 280 images). Images in the training set were used for model development and training whereas images in the validation set were used to optimize model hyperparameters, and finally, images in the test set were used for reporting the final performance and accuracy. Using the polygons, binary masks were generated for each image and then the images alongside their corresponding masks were separately split into smaller square image patches of different sizes (32 x 32, 64 x 64, 128 x 128, 256 x 256, and 512 x 512 pixels); this reduction in size was to improve the efficiency of model training. Among these patches, those that had at least one pixel annotated as CRS were then considered as positive samples for the classification task. The same number of patches were randomly selected among the images without CRS to keep the three image datasets (training, validation, and test) balanced.

#### 2.3.2 Image Analysis

Two different classes of machine learning models were applied to the data set to assess and describe CRS-affected foliage: classification, which determines the presence or absence of CRS for a given patch, and segmentation, which highlights the regions affected by CRS and quantifies the amount of CRS detected in the images. Six classification models were trained and evaluated with respect to their ability to classify patches containing plants exhibiting CRS symptoms. The classification models implemented were: ResNet18 (He et al. 2016); MobileNetV3 small, small custom, and large versions (Koonce 2021b); and EfficientNet-B3 and EfficientNet-B4 (Koonce 2021a). All of these models were initially trained and evaluated under the same conditions (same set of hyperparameters) on image patches of size 256 x 256 pixels and the best performing model was selected for further optimization over the range of patch sizes listed above. Finally, the top image patch classification model was used to approximately quantify the area of the affected regions of each image in the test set. Important to note that this was performed on the basis of image patches within the overall image and not on a per pixel basis. For image segmentation, three models were trained and evaluated including U-NET (Ronneberger, Fischer, and Brox 2015), Fully Convolutional Network (FCN) (Long, Shelhamer, and Darrell 2015), and DeepLabV3 (Chen et al. 2017). All models were trained and evaluated on image patch sizes of 256 x 256 pixels. Like the classification approach, the best performing model was then selected for further optimization over the range of patch sizes. Finally, the best performing models, with respect to classification and segmentation, were used to detect or quantify the regions, respectively, affected by CRS in the full-size images.

### 2.4 Model Training

#### 2.4.1 Model Evaluation

To evaluate performance during training and development of the classification models, binary cross entropy (BCE) was used as a loss function (Ho and Wookey 2020; Salehi, Erdogmus, and Gholipour 2017; Milletari, Navab, and Ahmadi 2016). The BCE represents a loss function utilized for binary classification, and uses the negative log of the error to penalize the model for incorrect predictions given by the following formula:

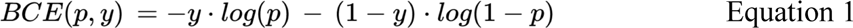

where *y* is the ground truth, integer label associated with the image and *p* is the prediction of the model, a numerical value ranging from zero to one. Note that the models output a value between zero and one for the prediction, and the penalty increases exponentially as the difference between prediction and the true label becomes greater.

To evaluate the performance of the segmentation models, the Dice coefficient was used (Ho and Wookey 2020; Salehi, Erdogmus, and Gholipour 2017; Milletari, Navab, and Ahmadi 2016). Dice coefficient was used as a loss function for semantic segmentation and represents a differentiable version of the intersection over union (IoU) metric. The IoU metric calculates the ratio between the predicted mask area and underlying ground truth data, which is the intersection between the expert-labeled masked area and the area predicted by the algorithm given the following formula:

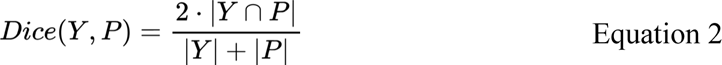

where *Y* is the ground truth mask and *P* is the predicted mask. The closer this value is to one, the more similar the prediction mask is to the ground truth.

The accuracy, precision, recall, and F1 score were computed for both classification and segmentation models to compare model performance. Accuracy values are computed as the ratio between the number of correct predictions and the total number of predictions and are provided for the classification and segmentation tasks. It is expressed using the number of true positives (TP), false positives (FP), true negatives (TN) and false negatives (FN) as follows:

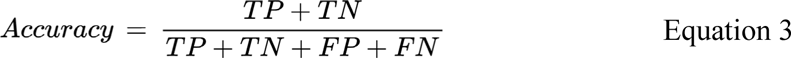

Accuracy is among the most widely used metric for classification and segmentation models, however, it has a significant limitation with respect to balance in the data set used. If the data set is unbalanced and the model incorrectly learns to only predict the majority label, accuracy merely reflects the class size ratio and nothing else. To provide additional model performance metrics that are independent of data set composition, the metrics of precision, recall and F1 score were used given the following formulas:

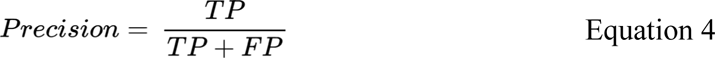

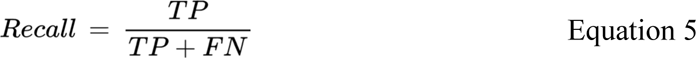

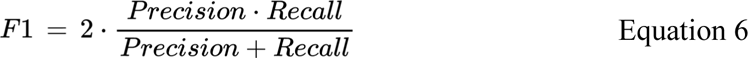

where *Precision* represents the portion of predicted values that are true positives, *Recall* represents the portion of true values that are correctly predicted by the model, and *F1* provides a harmonic mean of these two metrics. The combination of the above listed metrics provided a comprehensive assessment of the model performance during training, testing, and validation; however, *F1* was the metric used to select the best performing models.

Models were trained on two computing nodes: the first was an AMD EPYC 7542 32-Core processor (Advanced Micro Devices, Inc., Santa Clara, CA, USA), 1008 GB of RAM, and two NVIDIA A-100 40 GB GPU (NVIDIA Corporation, Santa Clara, CA, USA); and the second node was an Intel® Xeon® Gold 6146 CPU @ 3.20GHz 12-Core processor (Intel Corporation, Santa Clara, CA, USA), 188 GB of RAM, and an NVIDIA GeForce RTX 2080 Ti GPU (NVIDIA Corporation, Santa Clara, CA, USA). The learning rate was 1 x 10^−3^ and 3 x 10^−4^ for classification and segmentation tasks, respectively. A batch size of 32 with Adam optimizer was used for both classification and segmentation (Z. Zhang 2018). Models were trained for 50 epochs and used early stopping to avoid overfitting. Inference time was reported as the overall length of time from the model receiving an image patch to producing a segmentation or classification prediction.

#### 2.4.2 Evaluation of Image Patch Size on Accuracy and Processing Time

After the initial model evaluation experiments (i.e. training and evaluation of the classification and segmentation models on image patches sizes of 256 x 256 pixels) the best performing model with respect to F1 score on the validation data set were selected from both the classification and segmentation classes of models evaluated. These two models, one each from the classification and segmentation type, were then trained and evaluated on the datasets with different image patch sizes (32 x 32, 64 x 64, 128 x 128, 256 x 256, and 512 x 512 pixels) to assess the effect of image patch size on the model performance. Time benchmarking was performed on the aforementioned AMD EPYC server. In addition to the model performance on the test set, inference times were measured and reported. The code for training, inference, and evaluation of model performance can be accessed at: https://github.com/phytooracle/charcoal-dryrot-quantification.

## 3 RESULTS

### 3.1 Model Training

Nine models, including six classification and three segmentation, were trained to detect and segment the presence of CRS-affected sorghum plant tissue in images collected under field conditions (Figure 1). The two top-performing, EfficientNet-B3 and FCN, were able to distinguish and quantify, respectively, plant tissue that was exhibiting signs of CRS compared to plant tissue that was demonstrating the effects of drought stress. The ability to distinguish between the effects of CRS and drought stress is critical given that the pathology of CRS closely resembles that of drought stress making accurate discernments between the two challenging. Both algorithms were trained using a training data set consisting of 840 images, tested on 280 images, and validated using the remaining 280 images. The manually labeled images represented diverse field conditions, irrigation regimes, number of plants within an image, resolution, and image collection conditions creating a diverse representation of the pathology of CRS-impacted tissue with respect to real-world conditions that would be encountered either in a sorghum production or research setting. Several pre-processing steps were carried out during model training, which included transformations of color space and generation of image patches. The initial step involves converting the input images into grayscale color space. These grayscale images are utilized to create normalized image masks. Subsequently, these normalized image masks were used to generate small, non-overlapping patches that served as input for the model.

### 3.2 Model Deployment

The objective of the classification models is to categorize each image patch based on the presence or absence of CRS. If the model predicts the presence of CRS in a single patch, the image is labeled as CRS-affected. On the other hand, the segmentation models aim to classify each pixel within an image patch based on the presence or absence of CRS. The segmentation model quantifies the amount of CRS-affected pixels per patch and reports the sum of CRS-affected pixels for each image. The algorithm generates a results data file containing the image name and prediction results. For classification models, the prediction result is a binary “True” or “False” value for presence or absence of CRS, respectively. For segmentation models, the prediction results are the percentage of pixels classified as CRS compared to the total number of pixels in the image. For both classification and segmentation models, the total processing time for each image, which is the cumulative sum of the processing times for all individual patches within a single image, is also provided. Alongside the results data file, visual representations of the predictions are also produced. The predictions of CRS from the classification model are depicted in each patch of the full image (Figure 2A, left). In contrast, the segmentation model’s predictions are illustrated at the pixel level, emphasizing the specific regions of the leaves affected by CRS (Figure 2A, right). Users can select to run all classification and segmentation model classes tested here. In this case, the algorithm outputs a summary data file including the model name, patch size, and processing time for each each, similar to the results data file. The images for training, testing, and validation, both in their original and annotated forms, are available at: https://data.cyverse.org/dav-anon/iplant/projects/phytooracle/season_11_sorghum_yr_2020/level_0/charcol_rot_sorghum/dry_rot_raw.tar.gz. The code to train models and run inference are available on GitHub: https://github.com/phytooracle/charcoal-dryrot-quantification. To enhance the usability of these tools, the models have been incorporated into a Streamlit application that is publicly accessible: https://charcoal-dryrot-quantification.streamlit.app/. The application can also be executed locally by utilizing the associated Docker container: https://hub.docker.com/r/phytooracle/charcoal-dryrot-quantification. All models presented in the current study were integrated as a module in the PhytoOracle phenotyping workflow manager: https://github.com/phytooracle/automation/blob/main/yaml_files/other/crs_detection.yaml. The code and data are released under the open source MIT license.

### 3.3 Model Performance

#### 3.3.1 Assessing Model Performance and Identification of Top-Performing Models

Classification models were investigated to first assess whether CRS could be detected in images that also contained drought stressed sorghum plants given the symptoms of both stresses are visually similar although represent different classes (biotic and abiotic, respectively). Six classification models from three model families were selected for evaluation: ResNet18, MobileNet V3 using small, custom small, and large versions, and EfficientNet using B3 and B4 versions. All models were assessed on recall, accuracy, and F1 score, but F1 score was the main criterion for selection. The best performing model for the classification of image patches that contained at least one pixel of CRS-labeled plant tissue was EfficientNet-B3. This model had an accuracy of 86.97% and an F1 score of 0.73 (Table 1, Figure 2). The classification model with the lowest performance was ResNet18 with an accuracy of 82.62% and corresponding F1 score of 0.65. With respect to the other two classes of models tested, their average performance across models was as follows: MobileNet - accuracy of 84.99% and F1 score of 0.69; EfficientNet - accuracy of 86.44% and F1 Score of 0.72. EfficientNet-B3 had higher accuracy and F1 score so it was selected as the superior model (Table 2).

**Table 1.** Results of model training and testing on image patch sizes of 256 x 256 pixels for both tasks of classification and segmentation for different models in the respective categories. The loss function used for classification was binary cross entropy (BCE) whereas for segmentation, the dice coefficient was calculated. Models were developed using the training data set that consisted of 840 labeled images while the validation set consisted of 280 images. See text for description of performance metrics.

| Task/<br>Loss Function | Model Name | Validation<br>Accuracy | Validation<br>Precision | Validation<br>Recall | Validation<br>F1 |
| --- | --- | --- | --- | --- | --- |
| Classification/<br>BCE | ResNet18 | 82.62% | 0.75 | 0.61 | 0.65 |
|  | MobileNetV3 small | 85.47% | 0.78 | 0.68 | 0.70 |
|  | MobileNetV3 small custom | 84.65% | 0.78 | 0.66 | 0.69 |
|  | MobileNetV3 large | 84.84% | 0.77 | 0.64 | 0.68 |
|  | **EfficientNet-B3** | **86.97%** | **0.80** | **0.71** | **0.73** |
|  | EfficientNet-B4 | 85.91% | 0.78 | 0.68 | 0.71 |
| Segmentation/<br>Dice | U-NET | 97.49% | 0.69 | 0.62 | 0.62 |
|  | **FCN** | **97.76%** | **0.69** | **0.68** | **0.66** |
|  | DeepLabV3 | 97.93% | 0.72 | 0.65 | 0.65 |

**Table 2.** Impact of image patch size on model performance using the validation image set that consisted of 280 images. Performance metrics for EfficientNet-B3, a classification model, and Fully Convolutional Network (FCN), a segmentation model, with respect to the different size image patches, in pixels, used on the validation image set. Both models were trained on images that measured 256 x 256 pixels before being tested on the various image patch sizes. See text for description of performance metrics.

| *Model Name* | *Task* | *Patch Size* | *Validation Accuracy* | *Validation Precision* | *Validation Recall* | *Validation F1* |
| --- | --- | --- | --- | --- | --- | --- |
| *EfficientNet-B3* | *Classification* | 32 | 86.55% | 0.04 | 0.01 | 0.02 |
|  |  | 64 | 92.51% | 0.29 | 0.23 | 0.25 |
|  |  | 128 | 89.77% | 0.50 | 0.42 | 0.44 |
|  |  | 256 | 84.97% | 0.79 | 0.63 | 0.68 |
|  |  | **512** | **83.76%** | **0.89** | **0.83** | **0.84** |
| *FCN* | *Segmentation* | 32 | 95.13% | 0.16 | 0.13 | 0.14 |
|  |  | 64 | 95.57% | 0.22 | 0.15 | 0.17 |
|  |  | 128 | 97.12% | 0.43 | 0.38 | 0.39 |
|  |  | 256 | 97.80% | 0.71 | 0.65 | 0.65 |
|  |  | **512** | **98.48%** | **0.83** | **0.77** | **0.79** |

Given the results of the classification models and that CRS-affected tissue could be identified by machine learning algorithms, we next evaluated if the disease-affected regions could be segmented from the images. The expansion of our analyses to segmentation was driven by the question of whether desiccated tissue due to drought stress, which is similar in appearance to CRS-affected tissue, could reliably be distinguished. The three segmentation models evaluated were U-NET, FCN, and DeepLabV3 which were all trained on the same 840 labeled image data set as the classification models. With respect to model accuracy, both DeepLabV3 and FCN performed comparably with validation F1 scores of 0.65 and 0.66, respectively (Table 1). Of the two models, FCN had higher validation recall and F1 score, so it was selected as the superior model (Table 2). U-NET gave the lowest performance metrics with a validation F1 score of 0.62.

#### 3.3.2 Assessing the Impact of Image Patch Size on Model Performance

Next, we investigated the influence of image patch size on model performance to better understand how downstream applications, such as deployment on drone- or mobile-based phenotyping platforms, could be impacted by changes in image patch size. For both EfficientNet-B3 and FCN models run on the validation image set, there was a near linear increase in validation F1, recall, and precision scores as the image patch size increased from 32 x 32 pixels to the final size tested of 512 x 512 pixels (Table 2). One exception to this observed trend were the results validation accuracy. For the FCN model, the validation accuracy results increased from 95.13% to 98.48% for image patch sizes of 32 to 512 pixels, respectively. However, the EfficientNet-B3 exhibited contrasting results. The highest validation accuracy occurred for image patch size of 64 x 64 pixels and was 92.51%. For the following patch sizes of 128, 256, and 512 pixels, the validation accuracy decreased to 89.77%, 84.97%, and 83.76%, respectively.

#### 3.3.3 Assessing the Impact of Image Patch Size on Image Processing Time

In light of the impact that image patch size had on the performance metrics of the respective models, we next wanted to investigate how the patch sizes impacted image processing time as this is another significant factor with respect to algorithm deployment. Both models exhibited an exponential decrease in mean processing time with increasing image patch size. For the 32, 64, 128, 256, and 512 patch sizes the mean processing times were 263.34, 65.60, 24.85, 9.63, and 4.15 seconds and 518.09, 204.70, 70.52, 29.13, and 18.98 seconds for EfficientNet-B3 and FCN, respectively (Figure 3). Comparing the mean processing time within a model, we found that all processing times were significantly different (*p* < 0.05) for each image patch size level. Additionally, there was minimal variation observed for mean processing time for both models and image patch sizes; only FCN at 32 and 64 pixels exhibited any appreciable variation. With respect to the individual models themselves, it was not surprising to find that EfficientNet-B3, the classification model, was nearly twice as fast as the FCN, segmentation model, given the difference in tasks they perform.

**Figure 3.**
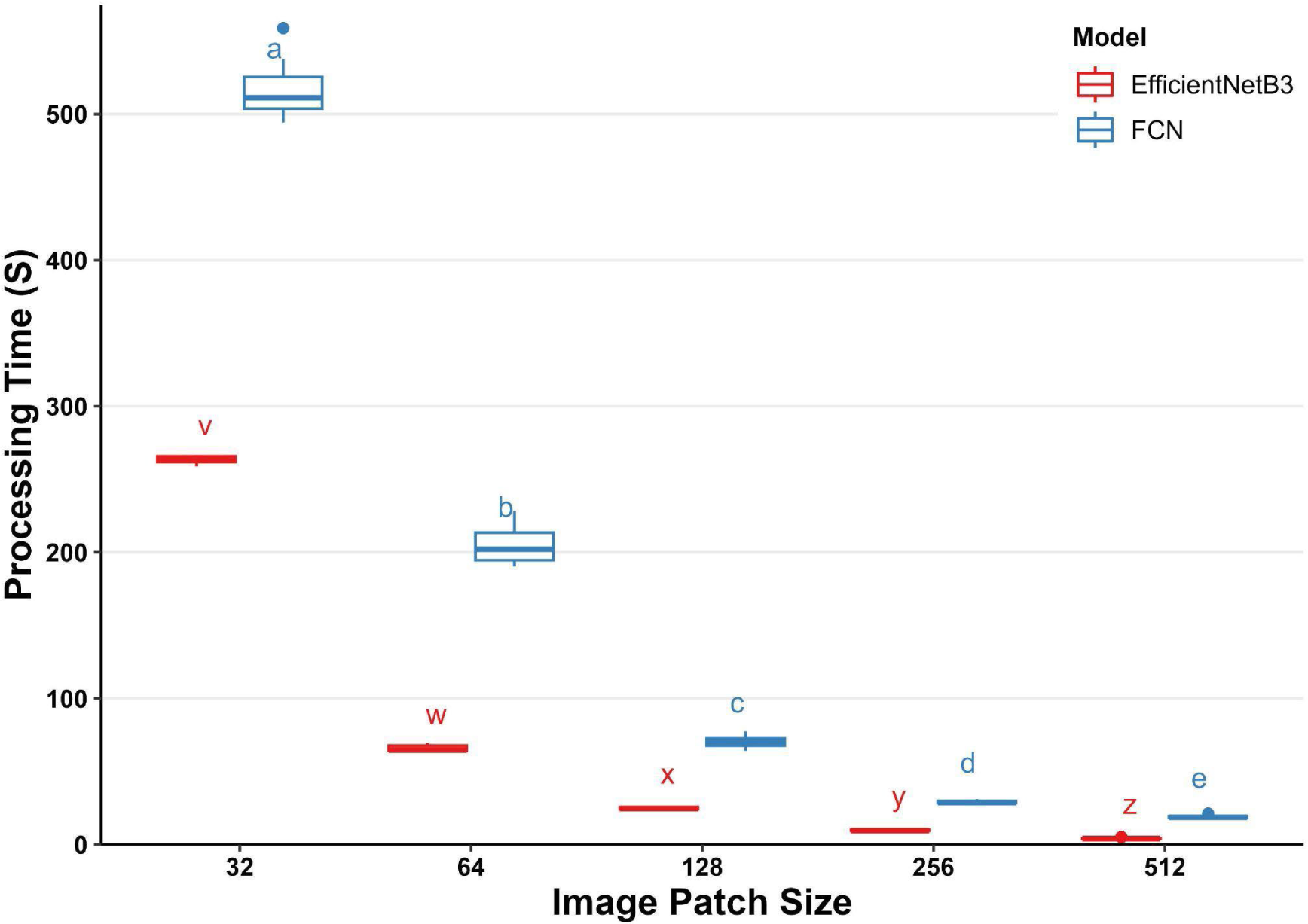
Relationship between image patch size and processing time in seconds (s) for the EfficientNet-B3 classification model and the Fully Convolutional Network (FCN) segmentation model. Different letters indicate significant differences among patch size processing times based on one-way analysis of variance (ANOVA) followed by Tukey’s post hoc test (p < 0.05). Error bars represent standard deviation.

#### 3.3.4 Evaluation of Model Performance on Test Set

With model training and validation completed, we next proceeded to evaluate how the respective models performed on the test data image set - the holdout image set. For the classification model EfficientNet-B3, the highest F1 score obtained was 0.70 for the 256 pixel image patch size. This is in contrast to the F1 score of 0.83 for the 512 pixel image patch size that was observed with the validation image set. With respect to accuracy, EfficientNet-B3 exhibited a near linear decrease in accuracy on the test image set with increasing patch size; however, the overall accuracy values were higher on the test image set compared to the validation image set (92.47% versus 87.51%, respectively). For the segmentation model FCN, the highest F1 score on the validation image set was 0.77 obtained on the 512 pixel image patch size; this also corresponded to an accuracy of 98.48%. On the test image set, the FCN model also obtained the largest F1 score on the 512 pixel image patch size - an F1 of 0.80 with a corresponding accuracy of 99.43%. With respect to all FCN performance metrics, the same trend was observed for both the validation and test image sets, namely that the metrics all increased with increasing image patch size. With respect to the image processing time, the same trends were observed for the test image set as for the validation set - an approximately exponential decrease in mean processing time per image as the image patch size increased. Additionally, and not surprisingly, the FCN model’s mean processing time was still much slower than the EfficientNet-B3 model. With respect to the individual models, the mean processing time for the image patches varied between the two models (Figure 3). The EfficientNet-B3 model had fairly consistent processing time per image with an average of 0.03 tenths of second whereas for the FCN model, the mean processing times for image patch sizes of 32, 64, 128, 256, and 512 were 0.02, 0.06, 0.08, 0.12, 0.35 tenths of second, respectively (Table 3).

**Table 3.** Performance metrics for each model are presented for the five different patch sizes to assess the effects of patch size on model performance and computational processing time. The performance metrics for EfficientNet-B3, a classification model, and Fully Convolutional Network (FCN), a segmentation model, with respect to the different size image patches (in pixels) and the time it took to process the whole image (seconds.tenths of second) as well as per image patch (seconds.tenths of second). Models were initially trained on image patch sizes of 256 x 256 pixels. See text for description of performance metrics.

| *Model Name* | *Task* | *Patch Size* | *Accuracy* | *Precision* | *Recall* | *F1* | *Mean Processing Time (image)* | *Mean Processing Time (patch)* |
| --- | --- | --- | --- | --- | --- | --- | --- | --- |
| *EfficientNet-B3* | *Classification* | 32 | 97.51% | 0.66 | 0.11 | 0.19 | 253.2 | 0.02 |
|  |  | 64 | 96.38% | 0.52 | 0.77 | 0.62 | 58.42 | 0.02 |
|  |  | 128 | 94.47% | 0.54 | 0.79 | 0.64 | 19.38 | 0.02 |
|  |  | **256** | **92.74%** | **0.64** | **0.77** | **0.70** | **6.42** | **0.03** |
|  |  | 512 | 81.24% | 0.53 | 0.88 | 0.66 | 2.27 | 0.05 |
| *FCN* | *Segmentation* | 32 | 97.90% | 0.37 | 0.69 | 0.48 | 311.93 | 0.02 |
|  |  | 64 | 98.81% | 0.58 | 0.56 | 0.57 | 192.15 | 0.06 |
|  |  | 128 | 98.99% | 0.61 | 0.79 | 0.69 | 65.96 | 0.08 |
|  |  | 256 | 99.19% | 0.67 | 0.83 | 0.74 | 25.95 | 0.12 |
|  |  | **512** | **99.43%** | **0.77** | **0.84** | **0.80** | **17.17** | **0.35** |

## 4 DISCUSSION

### 4.1 The Need of Phenotyping Tools for Detecting Biotic Stress

Charcoal rot of sorghum (CRS), a disease caused by the soilborne fungal pathogen *M. phaseolina*, poses a significant threat to sorghum, a critical grain crop. The pathogen, *M. phaseolina*, thrives in hot, dry conditions, which often coincide with the peak growing periods for sorghum in the southwest USA. As a result, areas with substantial sorghum production often experience high incidences of CRS, leading to considerable crop losses and economic impact (Marquez et al. 2021; Kaur et al. 2012). Effective management of CRS is therefore crucial to ensure the sustainability of sorghum production in these regions. Early detection of CRS is required for crop management systems for sorghum production and breeding improvement programs. As such, a major challenge is shifting CRS detection and quantification from manual visual assessment to automated, image-based phenotyping approaches. Handheld and drone-based RGB cameras provide a low cost solution, providing that there are accurate, precise, and efficient computer vision and AI/ML algorithms to process those data. This method not only allows for the detection and quantitative evaluation of disease severity but also supports the high-throughput screening of disease resistance variations in cultivars. Moreover, it acts as a crucial data gathering tool, potentially offering comprehensive insights into the prevalence, distribution, and effects of CRS across different environments.

To address this need, two classes of algorithms were evaluated for the classification and segmentation of CRS from RGB images of sorghum consisting of six and three base models of convolutional neural networks, respectively. The aim was to develop and evaluate models to cater to the diverse needs of sorghum researchers, breeders, and producers. The classification algorithm is more efficient, requiring fewer computational resources for deployment. This enables rapid assessment of CRS-affected tissues and is adaptable for use in the field on portable devices such as smartphones or drones. Users can efficiently identify CRS affected plants for near real-time decision making. On the other hand, the segmentation algorithm is more computationally intensive and currently requires dedicated computing resources. Users need to send images to off-field hardware to quantify the relative amount of above-ground affected plant tissues. To address this limitation, we developed a web application that can efficiently deploy both classification and segmentation models on multiple devices, including cell phones. Tools like these will enable breeders to select genotypes with improved resistance more effectively, potentially leading to increased future yield gains. Furthermore, these tools are directly applicable in crop management, as they can assess the severity of the disease and aid in planning treatment strategies such as control point spraying.

For CRS detection in images, binary classification can be applied on image patches. Alternatively, semantic segmentation can be used to quantify the ratio of pixels identified as CRS versus non-CRS. In this study, models representing each approach were trained and tested to evaluate performance metrics and assess suitability for the task of detecting and quantifying CRS. For classification, ResNet18 (He et al. 2016), MobileNetV3 small, custom small and large (Koonce 2021b) and EfficientNet-B3 and EfficientNet-B4 (Koonce 2021a) were implemented; for segmentation, U-NET (Ronneberger, Fischer, and Brox 2015), FCN (Long, Shelhamer, and Darrell 2015), and DeepLabV3 (Chen et al. 2017) were used. Each approach has advantages and disadvantages: classification models are generally faster, while segmentation models are often more accurate and have higher performance based on F1 score in particular.

### 4.2 Model Types for Plant Disease Detection and Quantification

#### 4.2.1 Classification Models

In ResNet18, stacks of convolutional layers alongside Max Pooling (Haibing Wu and Gu 2015), Batch Normalization (Ioffe and Szegedy 2015), and other auxiliary layers are combined in a feed forward network. The number 18 refers to the number of learnable layers. In addition, residual or skip connections are added between these blocks to (*i*) facilitate dealing with the vanishing gradient problem and (*ii*) to help the model learn more efficiently by having the capability of skipping some of the layers if necessary. Developed by Google (Mountain View, CA), MobileNetV3 models are designed for use on mobile devices and other embedded systems such as UAVs (Howard et al. 2017). These models are built out of depth-wise separable convolutions, a form of factorized convolutional layers that enables filters to be shared across channels, reducing the number of filters needed to improve computational efficiency. MobileNetV3 is the latest iteration of the MobileNet (Howard, Sandler, and Chu, n.d.). Small and Large MobileNets have a trade-off between latency and accuracy with the final layer of the MobileNetV3 Small and Large consisting of 576 and 1280 output neurons, respectively. To investigate the impact of reducing neuron count on performance, we customized MobileNetV3 Small by modifying the number of neurons in the last block of the model. This model has extra dense layers to reduce the number of output neurons more gradually from 576 to 288 to 64 to 1 neuron. (Tan and Le, n.d.) proposed a compound scaling method for scaling the depth, width, and resolution of neural network layers in a way to efficiently achieve better performance. They observed that while scaling up networks’ width, depth, and resolution generally improves accuracy because of higher network capacity, the dimensionality of the network needs to be adjusted properly in order to maximize these improvements on metrics while minimizing overfitting to the training data. In their paper, they propose a relationship between the scaling ratios for each dimension of the network which can be used to uniformly scale up the network to increase its efficiency and accuracy. Using their proposed relationship between the dimensions, they provide eight different architectures, namely EfficientNet-B0 - EfficientNet-B7. In this study we tested EfficientNet-B3 and EfficientNet-B4 based on the results presented in their paper.

#### 4.2.2 Segmentation Models

Fully convolutional neural networks (FCN) (Long, Shelhamer, and Darrell 2015) provide an end-to-end model for semantic segmentation of images with arbitrary sizes. Prior to FCNs, semantic segmentation used to be done by sliding a classification model over the entire image to get predictions for each pixel. U-NET is a fully convolutional neural network that has an encoder-decoder structure. It was developed by (Ronneberger, Fischer, and Brox 2015) to address the problem of semantic segmentation in biomedical images and uses skip connections. The encoder transforms the input image into a latent space with lower dimension and the decoder transforms the latent space into the output image, usually with the same dimension as the input. Our implementation varies slightly in architecture from the original paper where we used established best practices of adding padding to preserve input dimensions between downsampling and upsampling and added batch normalization between convolution layers and the Relu activation functions. DeepLabV3 (Chen et al. 2017) utilizes a novel method for enlarging the field of view of the convolution kernels to incorporate multi-scale context of images. This improves semantic segmentation results by using the atrous convolution (Holschneider et al. 1990).

### 4.3 Optimizing Model Performance and Addressing Limitations

#### 4.3.1 Effects of Image Patch Size on Performance and Processing Time

Processing time and model performance are closely linked to the size of image patches. When dealing with larger patches, the results are typically faster. This is likely due to larger patches containing more information, allowing the model to make decisions based on a broader context. This can be particularly beneficial for classification tasks, where the goal is to categorize the entire image or large regions of it. For instance, in crop disease detection, larger patches could help quickly identify whether a particular disease is present or absent in a field. On the other hand, smaller patches, while resulting in slower performance, can provide a more detailed assessment of the image. This can be especially useful when the goal is to assess the severity of a condition, such as the extent of disease in a crop field. Smaller patches allow for a more granular analysis, which can help quantify the extent of the disease and guide targeted interventions. Importantly, the choice of patch size in image analysis can significantly impact model performance, including the F1 score, a measure of a model’s accuracy. The F1 score considers both precision (how many selected items are relevant) and recall (how many relevant items are selected), making it a robust metric for model performance. In the case of EfficientNet-B3, a patch size of 512 was found to yield the highest validation F1 score (Table 3). This suggests that this patch size is optimal for balancing the trade-off between precision and recall in the model’s predictions. Interestingly, this patch size also has the fastest processing time per image (Figure 3). This means that not only does it provide accurate results, but it does so efficiently, making it a good choice for applications where both accuracy and speed are important. Similarly, for the FCN, a patch size of 512 results in the highest validation and test F1 score (Table 2, Table 3). This larger patch size allows the model to capture more contextual information, which can improve the accuracy of its predictions. Moreover, this patch size also has the fastest processing time per image, making it an efficient choice for image analysis tasks (Figure 3). Our results indicate that the choice of patch size can have a significant impact on the performance of image analysis models. Accuracy and processing speed are notable tradeoffs in the selection of patch size, which presents an opportunity for optimization of patch size to maximize these aspects. The choice of patch size in image analysis should be guided by the specific requirements of the task at hand and made on the basis of various performance metrics.

In the current study, EfficientNet-B3 and FCN performed best for the classification and segmentation of CRS, respectively (Table 1). As expected, processing time for a single image decreased as image patch size increased due to fewer patches being needed per image. While one may expect that increased patch size will result in decreased sensitivity, our results showed that the largest patch size tested for classification had the highest validation precision, recall, and F1 scores (Table 2). This is likely due to having more likelihood of dry rot being present in a larger patch. Therefore, the false positive rate would be lower compared to a smaller patch size, as we classify a prediction as a TP even if there’s only a single pixel of dry rot present in the patch. It should be noted that the validation accuracy was lowest for the largest patch size for classification.

Classification models work by assigning a single label to the entire image. These models are designed to focus on the most important features in the image that are relevant to the classification task. An increase in image patch size could introduce additional details and noise, such as background soil and neighboring plants, that may interfere with the main features, potentially leading to a decrease in validation accuracy. The increase in validation accuracy as the image patch size grows could be attributed to the additional context and detail that larger patches offer for the model to assess, thereby enhancing its performance.

#### 4.3.2 Limitations of Models

The results presented here increase the utility of these models for the classification and segmentation of CRS from image data, with the EfficientNet-B3 model taking ∼6.4 seconds to process an image with a patch-size of 256 x 256 pixels and the FCN model taking ∼17.2 seconds. These benchmarks utilized high-end GPUs, which are not currently deployable in the field. However, as GPU-based architectures continue to advance, the ability to run these models in the field using aerial-based or portable computing devices should be available in the near-term future, providing researchers, breeders, and growers the opportunity to detect and quantify CRS in real-time using automated crop phenotyping systems (Owens et al. 2008). Although existing models can be implemented on ground-based phenotyping systems equipped with powerful computers, the deployment of these models on UAVs is not yet feasible at scale. This limitation is primarily due to the intensive computational demands of the models, which surpass the hardware currently mountable on UAVs. Specifically, these models often need powerful GPUs to operate efficiently. The GPUs can perform parallel operations, making them ideal for the complex calculations and large data volumes involved in machine learning. However, the physical characteristics of GPUs, such as their size, weight, and power consumption, make them unsuitable for mounting on UAVs, which have strict limitations on payload capacity and power supply. Given the expected progress in hardware technology, it is reasonable to foresee that these models could be deployed on UAVs in the near future. It is therefore essential for research, like that presented here, to concentrate on the development, training, and refinement of these models to prepare them for widespread deployment on UAVs in the future. Another significant limitation of these models is the potential for ambiguity with other diseases or abiotic stresses. This means that the models may sometimes struggle to distinguish between the disease of interest and other similar-looking diseases or stress conditions. This could lead to false positives or negatives, impacting the accuracy of the model. The results presented here highlight the feasibility to discriminate heterogenous symptoms, namely drought and CRS, on a single leaf. Nonetheless, additional studies are required to determine whether this is applicable to other pairs of heterogeneous symptoms.

The quantification of disease tissue predicted in the image presents some challenges. The segmentation algorithm calculates the percentage of CRS relative to the entire image, not just the plant tissue. This approach was chosen because masking out the soil using vegetation indices could introduce errors, and, as a result, the reported result would be an amalgamation of various errors stemming from both the index calculation and the model prediction, complicating the interpretation and differentiation of these errors. The importance of having an unbiased and multi-dimensional system for quantifying plant disease instead of relying on human scoring is also worth noting. The process of human scoring can accumulate a variety of errors, particularly when multiple individuals are involved, as it can lead to subjectivity and inconsistency. An automated, unbiased system can provide more consistent and reliable results. Such a system could process large amounts of data quickly, making it a valuable tool for large-scale disease monitoring and management.

#### 4.3.3 The Importance of User-Friendly Phenotyping Tools

The importance of increased accessibility and integration of trained models cannot be overstated. By making these models easy to use, their use among non-experts increases. This democratization of technology is a key factor in driving innovation and progress in various fields. Integrating models into platforms like CyVerse, Streamlit, and PhytoOracle is a step towards this goal. These platforms are widely used and familiar to many users, making it easier for them to start using the models. Moreover, they provide an environment where users can immediately use the models and extract information, reducing the barriers to entry. Providing access to the code and models is another crucial aspect. This transparency not only allows users to understand how the models work but also enables them to modify and improve upon them. However, simply providing access to the code and models is not enough. It is equally important to provide integration into web applications and phenotyping workflow managers, such as PhytoOracle, to enable the computationally-efficient deployment of these models to large image datasets (Gonzalez et al. 2023). Increasing the accessibility and integration of training models, images, and results is of paramount importance; it empowers a wider range of users to leverage these models, fostering innovation and progress.

## 4 CONCLUSION

As the quantitative and qualitative results suggest, quantifying CRS in images using NNs is a difficult task because of the similarity between the symptoms of water deficit stress (ie. drought) and CRS. However, in this paper we proposed two approaches for this task using classification and segmentation models and we showed that quantifying CRS in plants exhibiting concurrent drought stress symptoms, despite being a difficult task, can be accomplished with a high level of accuracy. We showed that the segmentation models outperform the classification models in quantifying CRS. An extensive set of experiments were conducted to assess the effect of patch size on the processing time and performance of the models. We found that a patch size of 256 is suitable for classification and a patch size of 512 yields the best results for the segmentation models. The models were integrated into existing phenomics pipelines and a web application for user-friendly deployment of trained models on various platforms.

## ACKNOWLEDGMENTS

We would like to give our sincere gratitude to Cristian Salazar, Travis Simmons, Holly Ellingson, Victoria Ramsay, Brenda Esmeralda Jimenez, Hanna April Lawson, Hassan Alnamer, Jordan Pettiford, Michele Cosi, and Robert Strand for their assistance in image annotation and assistance with the field experiment.

## AUTHOR CONTRIBUTIONS

EG and AZ conceptualized and developed models, analyzed results, and wrote the manuscript. DR trained models and gathered results. SC, BR, and JD contributed to data analysis, image annotation, and image annotation review. CC contributed to data analysis. JH, AE, BD, KB, and EL contributed to project conceptualization, overseeing model development, and manuscript preparation. DP conceptualized, designed, and oversaw all aspects of the project including acquisition of funds and writing the manuscript. All authors contributed to the review of the manuscript and all authors have read the manuscript.

## DATA AVAILABILITY

The images used for training, validation, and testing the model can be accessed here: https://data.cyverse.org/dav-anon/iplant/projects/phytooracle/season_11_sorghum_yr_2020/level_0/charcol_rot_sorghum/dry_rot_raw.tar.gz. The code for model training and inference is available at: https://github.com/phytooracle/charcoal-dryrot-quantification. The models in this manuscript can be deployed on a Streamlit app at: https://charcoal-dryrot-quantification.streamlit.app/. The Docker container to run the Streamlit app locally is available at: https://hub.docker.com/r/phytooracle/charcoal-dryrot-quantification. The models were integrated as a module in the PhytoOracle phenotyping workflow manager: https://github.com/phytooracle/automation/blob/main/yaml_files/other/crs_detection.yaml.

## Abbreviations

AI: artificial intelligence
AZMET: Arizona Meteorological Network
BCE: binary cross entropy
CNN: convolutional neural network
CRF: conditional random field
CRS: charcoal rot of sorghum
EMS: ethyl methanesulfonate
FCN: fully convolutional network
FN: false negative
FP: false positive
GPU: graphics processing unit
IOU: intersection over union
ITS: internal transcribed spacer
MAC: Maricopa Agricultural Center
ML: machine learning
NLB: northern leaf blight of corn
NN: neural network
PDA: potato dextrose agar
RGB: red-green-blue
R-CNN: region-based convolutional neural network
SVWC: soil volumetric water content
TN: true negative
TP: true positive
UAV: unoccupied aerial vehicle
WL: water-limited
WW: well-watered

